# Elucidating syntrophic butyrate-degrading populations in anaerobic digesters using stable isotope-informed genome-resolved metagenomics

**DOI:** 10.1101/563387

**Authors:** Ryan M. Ziels, Masaru K. Nobu, Diana Z. Sousa

## Abstract

Linking the genomic content of uncultivated microbes to their metabolic functions remains a critical challenge in microbial ecology. Resolving this challenge has implications for improving our management of key microbial interactions in biotechnologies such as anaerobic digestion, which relies on slow-growing syntrophic and methanogenic communities to produce renewable methane from organic waste. In this study, we combined DNA stable isotope probing (SIP) with genome-centric metagenomics to recover the genomes of populations enriched in ^13^C after feeding ^13^C-labeled butyrate. Differential abundance analysis on recovered genomic bins across the SIP metagenomes identified two metagenome-assembled genomes (MAGs) that were significantly enriched in the heavy ^13^C DNA. Phylogenomic analysis assigned one MAG to the genus *Syntrophomonas*, and the other MAG to the genus *Methanothrix.* Metabolic reconstruction of the annotated genomes showed that the *Syntrophomonas* genome encoded all the enzymes for beta-oxidizing butyrate, as well as several mechanisms for interspecies electron transfer via electron transfer flavoproteins, hydrogenases, and formate dehydrogenases. The *Syntrophomonas* genome shared low average nucleotide identity (< 95%) with any cultured representative species, indicating it is a novel species that plays a significant role in syntrophic butyrate degradation within anaerobic digesters. The *Methanothrix* genome contained the complete pathway for aceticlastic methanogenesis, indicating that it was enriched in ^13^C from syntrophic acetate transfer. This study demonstrates the potential of stable-isotope-informed genome-resolved metagenomics to elucidate the nature of metabolic cooperation in slow-growing uncultured microbial populations, such as syntrophic bacteria and methanogens, that are important to waste treatment as well as global carbon cycling.

**Importance:** Predicting the metabolic potential and ecophysiology of mixed microbial communities remains a major challenge, especially for slow-growing anaerobes that are difficult to isolate. Unraveling the *in-situ* metabolic activities of uncultured species could enable a more descriptive framework to model substrate transformations by microbiomes, which has broad implications for advancing the fields of biotechnology, global biogeochemistry, and human health. Here, we investigated the *in-situ* function of mixed microbiomes by combining DNA-stable isotope probing with metagenomics to identify the genomes of active syntrophic populations converting butyrate, a C_4_ fatty acid, into methane within anaerobic digesters. This approach thus moves beyond the mere presence of metabolic genes to resolve ‘*who is doing what’* by obtaining confirmatory assimilation of labeled substrate into the DNA signature. Our findings provide a framework to further link the genomic identities of uncultured microbes with their ecological function within microbiomes driving many important biotechnological and global processes.

## Introduction

Linking microbial genomic identity with ecological function is considered a ‘Holy Grail’ in microbial ecology (1), and has broad implications for improving our ability to manage microbial communities in engineered biotechnologies. Anaerobic digestion is an example of a biotechnology that enables resource recovery from organic waste by generating methane gas as a renewable biofuel, and thus plays a role in establishing a circular economy (2). The production of methane in anaerobic digestion is executed through a series of trophic interactions constituting a metabolic network of fermenting bacteria, syntrophic acetogens, and methanogenic archaea (3, 4). Metabolic reconstructions based on shotgun metagenomic sequencing data have highlighted potential partitioning of functional guilds within anaerobic digester microbiomes (4). Yet, our understanding of the ecophysiology of the microorganisms present in anaerobic digesters is limited by the high community complexity and lack of cultured representatives (4). Elucidating the nature of interspecies interactions between different trophic groups in the anaerobic digester metabolic network could help to better understand and optimize the conversion of organic wastes into renewable methane.

The terminal steps in the anaerobic metabolic network — syntrophy and methanogenesis — are considered rate limiting steps for the production of methane from organic substrates (5). The syntrophic oxidation of fatty acids is also responsible for a considerable portion of carbon flux in methanogenic bioreactors, as fatty acids are often produced during fermentation of mixed organic substrates (6). The accumulation of fatty acids in anaerobic digesters is often responsible for a reduction in pH and process instability (3). In particular, syntrophic degradation of the 4-carbon fatty acid, butyrate, can be a bottleneck for anaerobic carbon conversion, as this metabolism occurs at the thermodynamic extreme. Butyrate degradation to acetate and hydrogen is thermodynamically unfavorable under standard conditions (ΔG°’= 53 kJ/mol), and only yields - 21 kJ/mol under environmental conditions typical of anaerobic bioreactors (pH 7, 1 mM butyrate and acetate, 1 Pa H_2_). Thus, cooperation with acetate-and hydrogen-scavenging methanogenic partners is necessary to maintain thermodynamic favorability. Cultured representative species carrying out syntrophic fatty acid oxidation are potentially underrepresented due to their slow cell yields and difficulty of isolation in the lab (7). Insofar, only two mesophilic (*Syntrophomonas* and *Syntrophus*) and two thermophilic (*Syntrophothermus* and *Thermosyntropha*) genera (12 bacterial species total) have been shown to oxidize butyrate in syntrophic cooperation with methanogenic archaea, and they all belong to the families *Syntrophomonadaceae* and *Syntrophaceae* (7). Despite their major roles in processing carbon within anaerobic bioreactors, many syntrophic fatty acid-oxidizing bacteria have evaded detection with quantitative hybridization-based techniques (8), which is likely due to their low biomass yields (9) or our incomplete knowledge of active syntrophic populations within anaerobic digesters (10). Broad metagenomic surveys of anaerobic digester communities have similarly observed poor resolution of syntrophic populations, owing to their low abundance (4, 11). Thus, highly sensitive culture-independent approaches are needed to expand our understanding of the ecophysiology of syntrophic populations to better control and predict metabolic fluxes in anaerobic environments.

Recently, we demonstrated the potential of combining DNA-stable isotope probing (SIP) with genome-resolved metagenomics to identify syntrophic populations degrading the long-chain fatty acid, oleate (C_18:1_), within anaerobic digesters (12). Stable-isotope informed metagenomic sequencing can enrich metagenomic libraries with genomic sequences of actively-growing microbes that incorporate ^13^C into their biomass from an added labeled substrate (13), and thus allows for a ‘zoomed in’ genomic view of low-abundance populations such as syntrophs. We also demonstrated that this approach was amenable for recovering high-quality microbial genomes using a differential-coverage based binning approach, as genomes from active microbes have low abundance in DNA from ^12^C controls but are enriched in ^13^C-ammended treatments (12). Here, we applied stable-isotope informed metagenomics to resolve the genomic makeup of active syntrophic butyrate-degrading populations within an anaerobic digester. We utilized biomass collected from the same anaerobic digesters as were previously used for DNA-SIP with oleate (12) at a similar time point, thus allowing for genomic comparisons using a multi-substrate SIP dataset. This approach identified potential metabolic flexibility in syntrophic populations processing multiple fatty acids within the anaerobic digesters, and elucidated the genomic identity of syntrophic partnerships between active methanogens and bacteria.

## Results and Discussion

### DNA stable isotope probing (SIP) of methanogenic microcosms with ^13^C-labeled butyrate

Anaerobic digester contents from a pulse-fed and continuous-fed anaerobic digester were incubated in duplicate microcosms that were spiked with either ^12^C or ^13^C-labelled butyrate (40 mM) for approximately 50 hours. The added butyrate was converted into methane at >80% conversion efficiency based on COD recovery (Supplemental Figure S1). After the 50 hr incubation, the contents of the microcosms were sacrificed for DNA extraction, density-gradient centrifugation, and fractionation.

The abundance of 16S rRNA genes of the known butyrate-degrading genus, *Syntrophomonas*, was quantified across density gradient fractions using qPCR to identify DNA fractions that were enriched in ^13^C (Supplemental Figure S2). Density fractions with a buoyant density from 1.70 to 1.705 had 2.0 to 2.2-times higher *Syntrophomonas* 16S rRNA genes (normalized to maximum concentration) than the ^12^C controls. Those DNA fractions were selected from each SIP microcosm for metagenomic sequencing, as well as for 16S rRNA gene amplicon sequencing.

The microbial communities in the heavy density-gradient fractions were assessed through paired-end 16S rRNA gene amplicon sequencing for all ^12^C-and ^13^C-incubated duplicate microcosms (Figure 1). Differential abundance analysis of OTU read counts with DESeq2 (14) showed that approximately 50% (7 of 15) of the significantly-enriched (*p<*0.05) OTUs in the ^13^C heavy DNA samples relative to ^12^C were taxonomically classified as *Syntrophomonas* for the pulse-fed digester (Supplemental Figure S3). For the continuous-fed digester, approximately 40% of the ^13^C-enriched OTUs (7 of 17) were assigned to *Syntrophomonas* (Supplemental Figure S4). Additionally, two ^13^C-enriched OTUs in both digesters were assigned to *Methanothrix* (formerly *Methanosaeta*), which likely scavenge the ^13^C-acetate generated by *Syntrophomonas* during ^13^C-butyrate degradation. While one previous study observed that *Syntrophaceae* was predominantly enriched in anaerobic digester granular sludge incubated with ^13^C-labeled butyrate (10), various other studies also detected *Syntrophomonadaceae* populations as active syntrophic butyrate degraders in anaerobic digester sludge using ^14^C-labeled butyrate and MAR-FISH (15), in anaerobic digester sludge through SIP using ^13^C-labeled oleate (12), and in rice paddy soil with SIP using ^13^C-labeled butyrate (16). In the latter two studies, acetate-scavenging partners (*Methanothrix* and *Methanosarcinaceae*) were also enriched. Indeed, syntrophic interaction with acetoclastic methanogens is beneficial as acetate accumulation can thermodynamically hinder butyrate oxidation (*e.g.*, ∆G exceeds the theoretical threshold for catabolism [-10 kJ/mol] when acetate accumulates beyond 10 mM [pH 7, 1mM butyrate, and 1Pa H_2_]). Notably, H_2_-and formate-consuming methanogens necessary for syntrophy are not detected during degradation of ^13^C butyrate because these archaea utilize CO_2_ as a carbon source.

**Figure 1:**
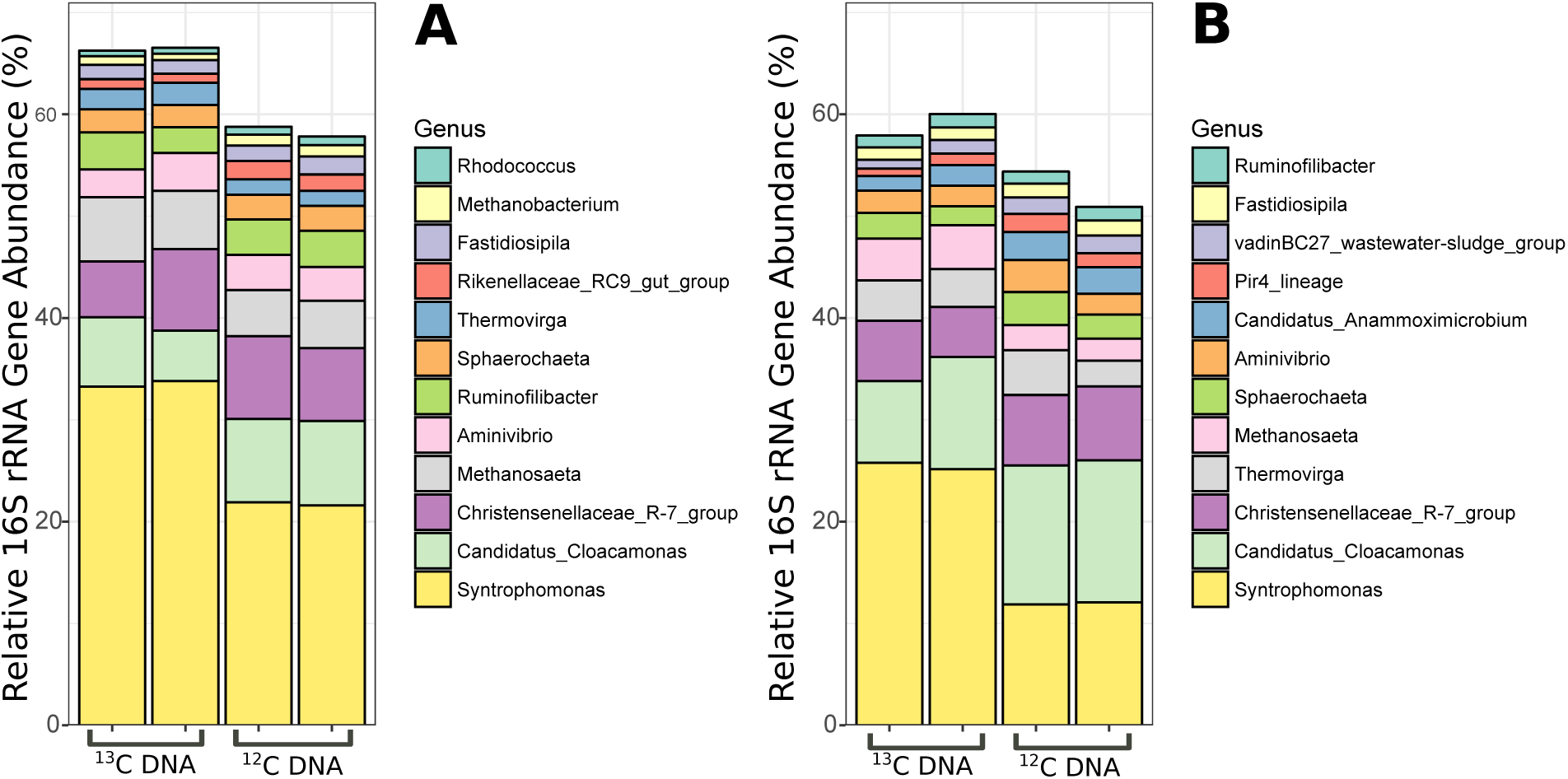
Relative 16S rRNA gene amplicon sequence abundance of the top 12 most abundant prokaryotic genera in the ^13^C butyrate-amended microcosms and the ^12^C butyrate-amended controls for (A) the pulse-fed codigester and (B) the continuous-fed codigester.

Our results also found ^13^C-enriched OTUs from lineages not known to degrade butyrate under methanogenic conditions: *Treponema, Luteimonas, Thauera, Christensenellaceae* (*Firmicutes*), and *Anaerolineaceae* (*Chloroflexi*) (Supplemental Figures 3 and 4). Other studies using ^13^C-butyrate also detected enrichment of populations likely unable to degrade butyrate, including *Tepidanaerobacter* and *Clostridium* in a thermophilic anaerobic digester operated at 55ºC (10) and *Chloroflexi* and *Planctomycetes* in rice paddy soil (16). Members of *Tepidanaerobacter* and *Clostridium* are known to syntrophically oxidize acetate under thermophilic conditions (17), and may have thus been enriched in ^13^C RNA from ^13^C-labeled acetate produced during the beta-oxidation of labeled butyrate in the study by Hatamoto *et al.* (10). Similarly, the *Chloroflexi* and *Planctomycetes* populations were hypothesized to have become enriched due to cross-feeding of intermediate metabolites like acetate in the rice paddy soil (16). Thus, the ‘peripheral’ populations detected in our study may grow on cell-decay products, as genome-resolved metagenomics recently indicated that some uncultured *Anaerolineaceae* species are likely fermenters in anaerobic digesters (18). These results thus suggest that carbon cross-feeding may occur between multiple microbial groups during the syntrophic degradation of butyrate in anaerobic digesters.

### Identifying active metagenome-assembled genomes (MAGs) in SIP metagenomes

Metagenomic sequencing of heavy DNA from duplicate ^13^C and ^12^C-butyrate amended microcosms yielded an average of 30 M paired reads per sample for both digesters (*n*=8) (Supplemental Table S1). The filtered reads from heavy ^13^C DNA were co-assembled, yielding a total assembly length of 516 Mb of contigs larger than 1 kb, with an average (N50) contig length of 5 kb. The fraction of filtered short reads that mapped to the co-assembly were 66% ± 3 (s.d) and 69% ± 1 for the ^12^C and ^13^C metagenomes, respectively (*n*=4 each) (Supplemental Table S1). The co-assembly generated from ^13^C reads thus captured much of the genomic information present in the heavy DNA fractions.

The assembled metagenomic contigs were organized into 160 genomic bins at various levels of completion and redundancy (Supplemental File 1). Differential abundance analysis of the mapped read counts for the bins across the ^13^C and ^12^C metagenomes with DESeq2 (14) identified two genomic bins that were significantly (*p* < 0.05) enriched in ^13^C DNA (Table 1). Based on suggested completion and redundancy metrics for metagenome-assembled-genomes (MAGs) (19), one genomic bin is classified as a high-quality MAG (completion >90%, redundancy <10%), while the other is a medium-quality MAG (completion >50%, redundancy <10%). Taxonomic classification with CheckM (20) assigned one of the MAGs to the genus *Syntrophomonas*, and the other to *Methanothrix* (Table 1).

**Table 1:** Genomic feature summary of the two metagenome-assembled genomes that were significantly enriched in ^13^C after the degradation of labeled butyrate.

| Name | Bin ID | Taxonomy<sup>1</sup> | Size (Mb) | GC (%) | Completion (%)<sup>2</sup> | Redundancy (%)<sup>2</sup> | Quality Classification<sup>3</sup> |
| --- | --- | --- | --- | --- | --- | --- | --- |
| *Syntrophomonas* BUT1 | Bin 26_1 | *Syntrophomonas* | 2.87 | 51.2 | 96.4 | 1.4 | High Quality Draft |
| *Methanotherix* BUT2 | Bin 26_2 | *Methanotherix* | 1.44 | 53.6 | 74.7 | 3.1 | Medium Quality Draft |

<sup>1</sup> Based on phylogenetic placement of single marker genes with CheckM (72)
<sup>2</sup> Measured with Anvi'o (72)
<sup>3</sup> Based on parameters suggested by Bowers *et al.* (19)

Phylogenomic placement of the ^13^C-enriched *Syntrophomonas* BUT1 MAG was consistent with its taxonomic assignment, as it was located in the *Syntrophomonas* genome cluster within the family *Syntrophomonadaceae* (Figure 2). The closest relative to *Syntrophomonas* BUT1 based on single-copy marker genes was *Syntrophomonas* PF07, which was a genomic bin enriched in ^13^C from DNA-SIP with labelled oleate (^13^C_18:1_) with sludge from the same pulse-fed digester used in this study (12). A high average nucleotide identity (ANI) of 99% was observed between the *Syntrophomonas* BUT1 and *Syntrophomonas* PF07 genomes (Supplemental Figure S5), suggesting that these two organisms likely originated from the same sequence-discrete population (21). The next closest relative of *Syntrophomonas* BUT1 based on the phylogenomic analysis was *Syntrophomonas zehnderi* OL-4 (Figure 2), which was isolated from an oleate-fed anaerobic granular sludge bioreactor (22). However, the ANI between *Syntrophomonas* BUT1 and *Syntrophomonas zehnderi* OL-4 was below 95% (Supplemental Figure S5), suggesting that these two organisms were different species (23). Thus, the active butyrate-degrading bacterial MAG identified in this study is distinct from any species obtained in isolation at this time. The detection of the sequence-discrete population of *Syntrophomonas* BUT1 within heavy ^13^C-DNA from experiments with both labelled butyrate and oleate indicate that this syntrophic population could be metabolically flexible; that is, it may grow on fatty-acids of variable length and degree of saturation. This finding has implications for current frameworks for mathematical modeling of anaerobic digesters, which typically assume that LCFA and butyrate-degrading populations are distinct (24). Thus, the incorporation of genomic and functional characterization, as obtained through DNA-SIP genome-resolved metagenomics, may help to improve our ability to accurately model anaerobic digestion processes by accounting for metabolic flexibility within key functional guilds.

**Figure 2:**
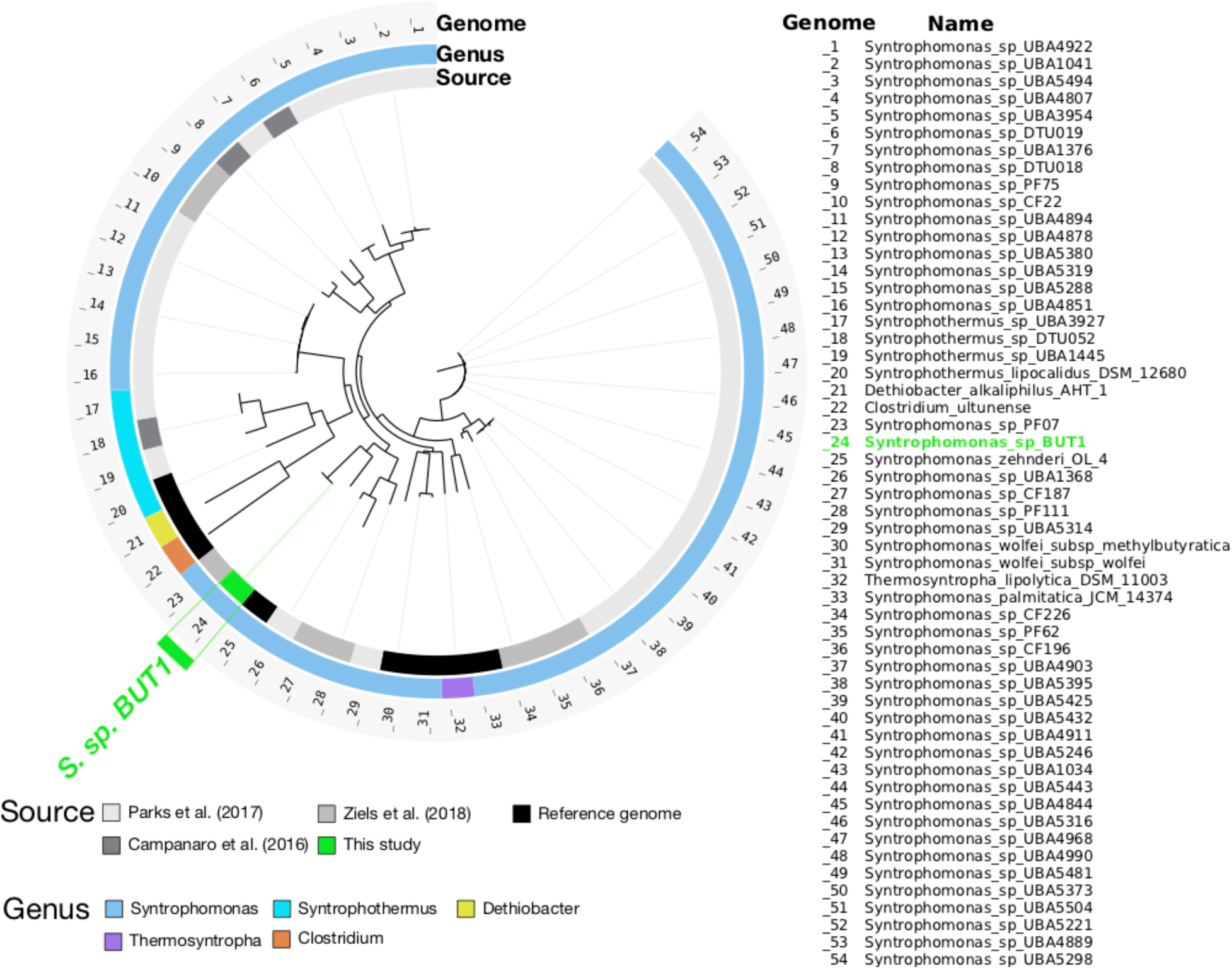
A phylogenomic tree showing the relationship of the ^13^C-enriched *Syntrophomonas* BUT1 to other genomes available from the *Syntrophomonadaceae* family in the NCBI nr database (downloaded April, 2018). The tree is based on a concatenated alignment of 139 bacterial single copy marker genes (75) obtained using Anvi’o (72). Open reading frames were predicted with Prodigal v.2.6.3 (69), and queried against database of bacterial and archaeal single copy marker genes using HMMER v.2.3.2 (76). The tree was calculated using FastTree (77). The *Clostridium ultunense* genome was used as the outgroup.

A phylogenomic analysis of the ^13^C-enriched *Methanothrix* BUT2 based on archaeal single-copy marker genes placed the MAG within the genus *Methanothrix,* consistent with its taxonomic assignment (Figure 3). *Methanothrix* BUT2 was closely clustered with the genome of *Methanothrix soehngenii* GP6, along with four MAGs reported in the study of Parks *et al.* (25). Congruent with the phylogenomic analysis, *Methanothrix* BUT2 shared an ANI over 98% with *Methanothrix soehngenii* GP6 and the same with four MAGs from Parks *et al.* (25) (*M.* UBA243, *M.* UBA458, *M.* UBA70, *M.* UBA356), indicating that these genomes likely form a sequence-discrete population (Supplemental Figure S5). A second, closely-related population including three MAGs from Parks *et al.* (25) (*M.* UBA372, *M.* UBA332, *M.* UBA553) shared an ANI of 96% with the *Methanothrix* BUT2 population (Supplemental Figure S5).

**Figure 3:**
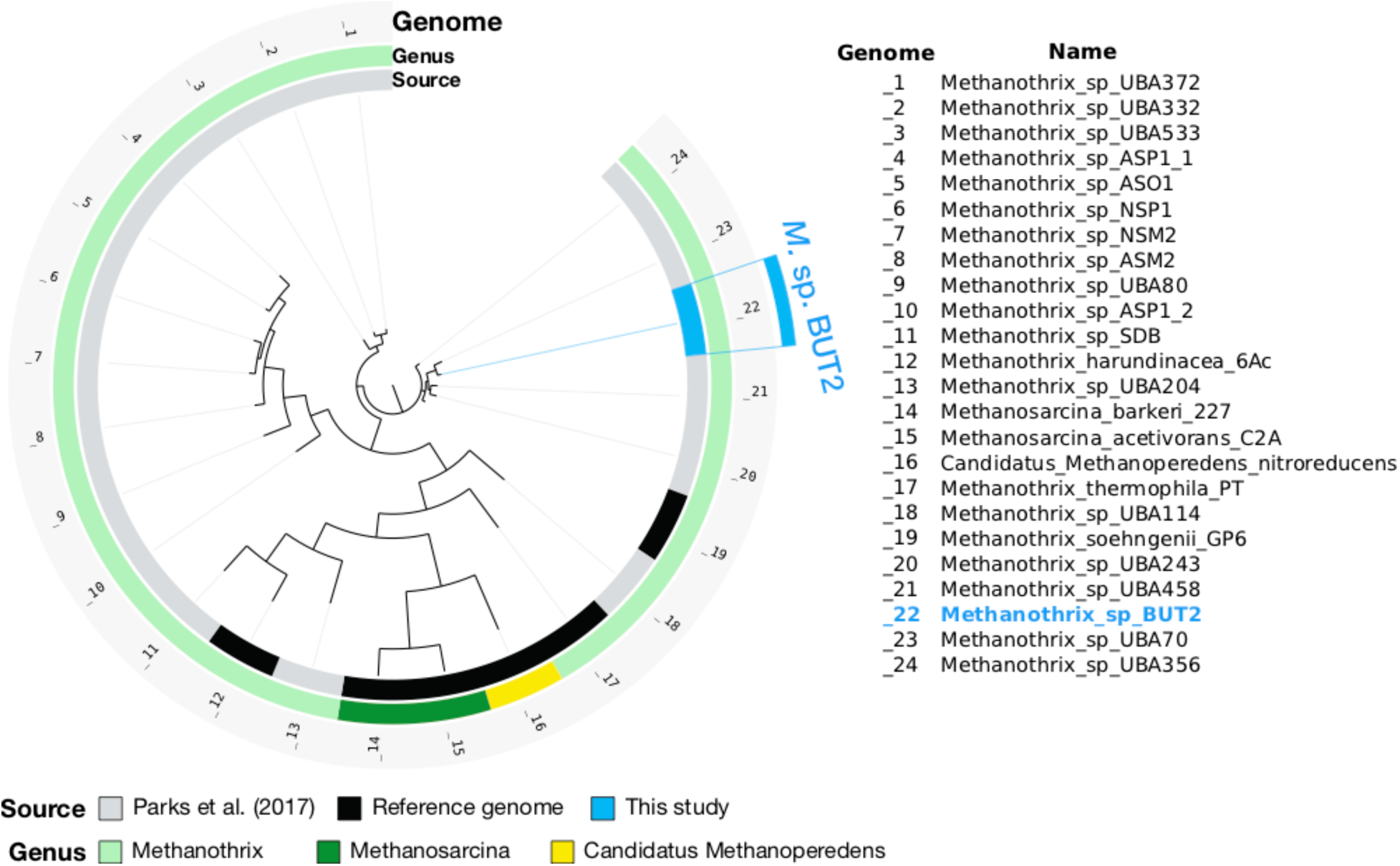
A phylogenomic tree showing the relationship of the ^13^C-enriched *Methanothrix* BUT2 to other genomes within the order *Methanosarcinales* in the NCBI nr database (downloaded April, 2018). The tree is based on a concatenated alignment of 162 archaeal single copy marker genes (78) obtained using Anvi’o (72). Open reading frames were predicted with Prodigal v.2.6.3 (69), and queried against database of bacterial and archaeal single copy marker genes using HMMER v.2.3.2 (76). The tree was calculated using FastTree (77). The *Candidatus* Methanoperedens nitroreducens genome was used as the outgroup.

DNA-SIP using ^13^C-labeled oleate with the same anaerobic digester biomass as this study did not identify any ^13^C-enriched methanogenic archaea in the genome-resolved metagenomic analysis (12). One possible explanation for the higher relative enrichment of methanogens on ^13^C-butyrate versus ^13^C-oleate could be the higher fraction of overall free-energy partitioned towards methanogens during anaerobic butyrate degradation versus oleate degradation. For the overall conversion of 1 mole of butyrate to CO_2_ and CH_4_ at environmental conditions in anaerobic digesters, the thermodynamic yields would be −21.1, −9.4, and −58.9 kJ for the acetogenic bacteria, hydrogenotrophic methanogens, and aceticlastic methanogens, respectively (Table 2). For similar conversion of 1 mole of oleate, the thermodynamic yields would be −219.9, −70.6, and −264.9 kJ respectively (Table 2). Thus, the acetogen would gain a much lower percentage of the overall free energy yield from conversion of butyrate (24%) than oleate (40%). As cell yield can depend on free-energy (26), the lower yield of the butyrate degradation would likely leave a higher fraction of acetate for assimilation by aceticlastic methanogen compared to oleate. Supporting this, the relative energy yield of aceticlastic methanogens compared to the acetogen is higher for conversion of butyrate (0.45) than oleate (0.32). As the stable-isotope informed analysis utilized in this study depended on heterotrophic incorporation of the added ^13^C into biomass, it was not expected that autotrophic (i.e. hydrogenotrophic) methanogens would be enriched in the heavy ^13^C DNA because no CO_2_ is produced during butyrate beta-oxidation (Table 2). Comparing the enriched communities from DNA SIP with different fatty acids, along with bicarbonate, could potentially highlight differences in energy partitioning between syntrophic bacteria and different archaeal partners.

**Table 2:**
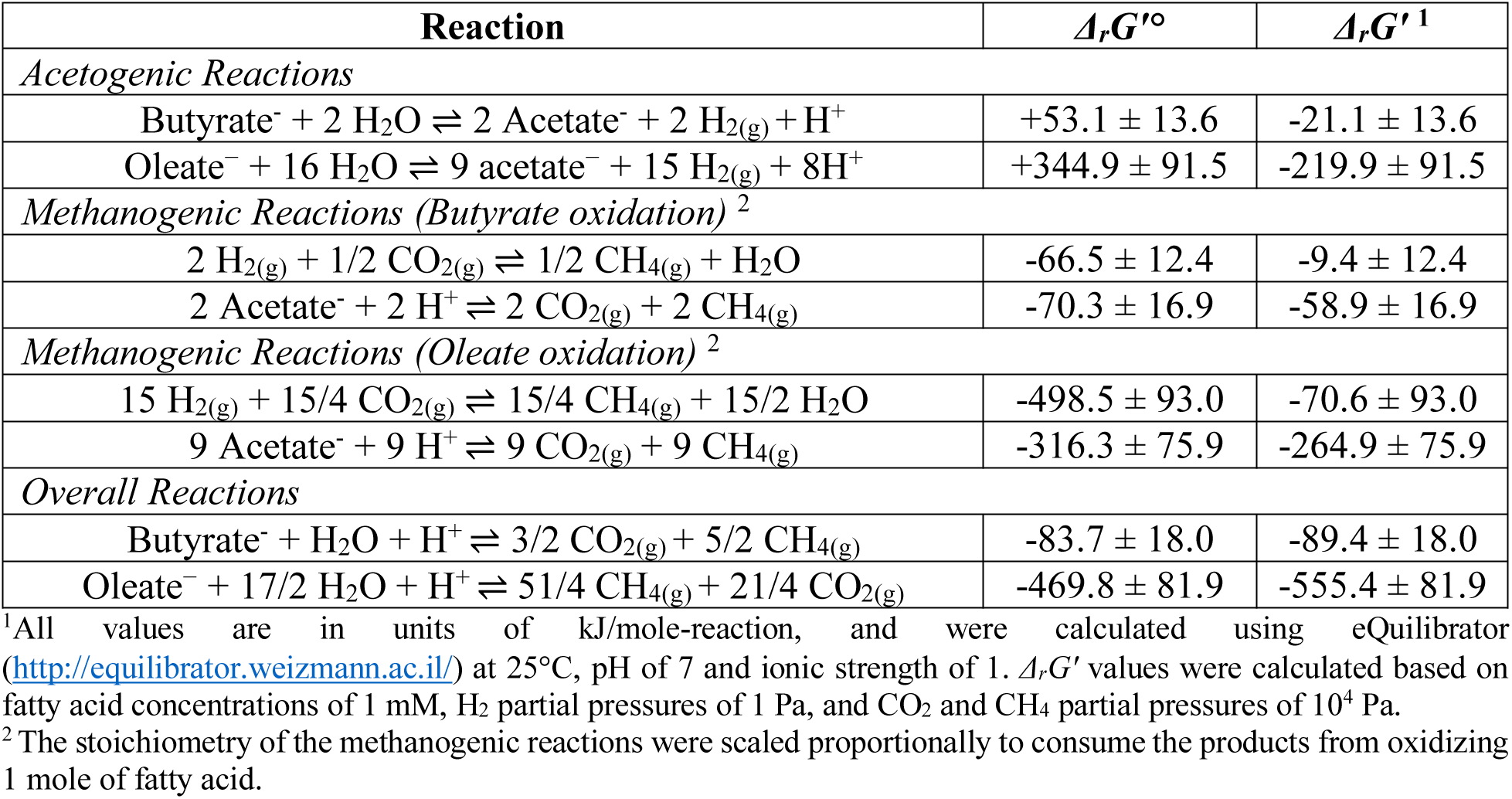
Gibbs free energy for some of the acetogenic and methanogenic reactions likely involved in the syntrophic conversion of butyrate and oleate.

| Reaction | $\Delta_r G'^{\circ}$ | $\Delta_r G'^1$ |
| --- | --- | --- |
| *Acetogenic Reactions* |  |  |
| $\text{Butyrate}^- + 2 \text{H}_2\text{O} \rightleftharpoons 2 \text{Acetate}^- + 2 \text{H}_{2(\text{g})} + \text{H}^+$ | $+53.1 \pm 13.6$ | $-21.1 \pm 13.6$ |
| $\text{Oleate}^- + 16 \text{H}_2\text{O} \rightleftharpoons 9 \text{acetate}^- + 15 \text{H}_{2(\text{g})} + 8 \text{H}^+$ | $+344.9 \pm 91.5$ | $-219.9 \pm 91.5$ |
| *Methanogenic Reactions (Butyrate oxidation)<sup>2</sup>* |  |  |
| $2 \text{H}_{2(\text{g})} + 1/2 \text{CO}_{2(\text{g})} \rightleftharpoons 1/2 \text{CH}_{4(\text{g})} + \text{H}_2\text{O}$ | $-66.5 \pm 12.4$ | $-9.4 \pm 12.4$ |
| $2 \text{Acetate}^- + 2 \text{H}^+ \rightleftharpoons 2 \text{CO}_{2(\text{g})} + 2 \text{CH}_{4(\text{g})}$ | $-70.3 \pm 16.9$ | $-58.9 \pm 16.9$ |
| *Methanogenic Reactions (Oleate oxidation)<sup>2</sup>* |  |  |
| $15 \text{H}_{2(\text{g})} + 15/4 \text{CO}_{2(\text{g})} \rightleftharpoons 15/4 \text{CH}_{4(\text{g})} + 15/2 \text{H}_2\text{O}$ | $-498.5 \pm 93.0$ | $-70.6 \pm 93.0$ |
| $9 \text{Acetate}^- + 9 \text{H}^+ \rightleftharpoons 9 \text{CO}_{2(\text{g})} + 9 \text{CH}_{4(\text{g})}$ | $-316.3 \pm 75.9$ | $-264.9 \pm 75.9$ |
| *Overall Reactions* |  |  |
| $\text{Butyrate}^- + \text{H}_2\text{O} + \text{H}^+ \rightleftharpoons 3/2 \text{CO}_{2(\text{g})} + 5/2 \text{CH}_{4(\text{g})}$ | $-83.7 \pm 18.0$ | $-89.4 \pm 18.0$ |
| $\text{Oleate}^- + 17/2 \text{H}_2\text{O} + \text{H}^+ \rightleftharpoons 51/4 \text{CH}_{4(\text{g})} + 21/4 \text{CO}_{2(\text{g})}$ | $-469.8 \pm 81.9$ | $-555.4 \pm 81.9$ |

<sup>1</sup>All values are in units of kJ/mole-reaction, and were calculated using eQuilibrator (http://equilibrator.weizmann.ac.il/) at 25°C, pH of 7 and ionic strength of 1. $\Delta_r G'$ values were calculated based on fatty acid concentrations of 1 mM, $\text{H}_2$ partial pressures of 1 Pa, and $\text{CO}_2$ and $\text{CH}_4$ partial pressures of $10^4$ Pa.
<sup>2</sup>The stoichiometry of the methanogenic reactions were scaled proportionally to consume the products from oxidizing 1 mole of fatty acid.

### Metabolic potential of ^13^C-enriched MAGs

Functional annotation and metabolic reconstruction of the ^13^C-enriched MAGs revealed their capacity to metabolize the labeled butyrate into methane through syntrophic cooperation (Figure 4).

**Figure 4:**
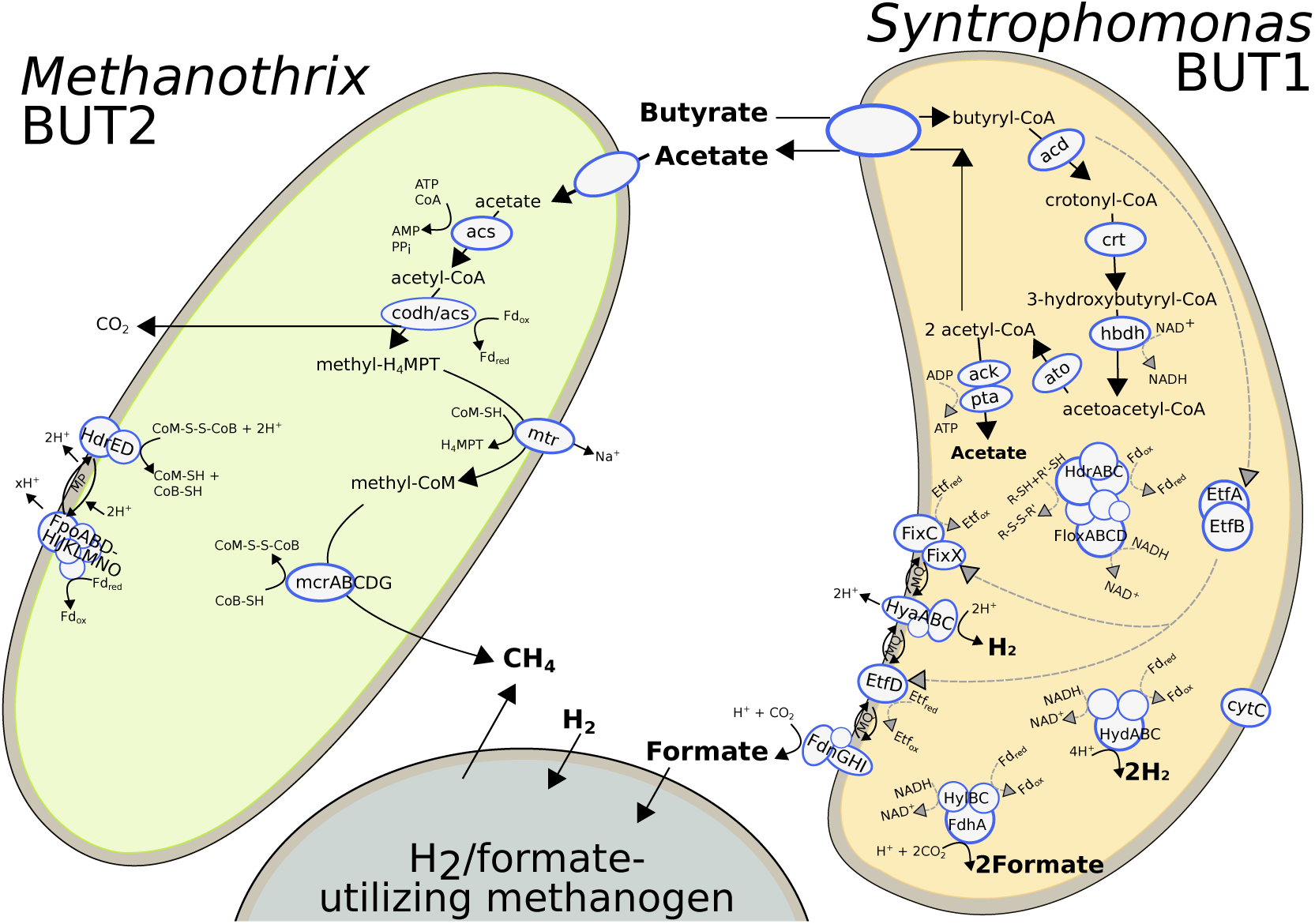
Cell diagram showing detected metabolic pathways for anaerobic butyrate degradation in syntrophic cooperation between *Syntrophomonas* BUT1 and *Methanothrix* BUT2. Abbreviations of enzymes are defined in Supplemental Files 2 and 3. The H_2_/formate utilizing methanogenic partner is shown for conceptual purposes, but was not identified with ^13^C-DNA SIP this study due to their autotrophic growth in the microcosms.

A complete pathway for butyrate β-oxidation was annotated in *Syntrophomonas* BUT1, indicating that this MAG was capable of metabolizing the added ^13^C-butyrate (Figure 4). Notably, several homologues were detected for genes in the β-oxidation pathway (Supplemental File 2). *S.* BUT1 genome encodes 6 acyl-CoA-transferases, 7 acyl-CoA dehydrogenases, 8 enoyl-CoA hydratases, 5 3-hydroxybutyryl-CoA dehydrogenases, and 10 acetyl-CoA acetyltransferases (Supplemental File 2). The presence of homologous β-oxidizing genes was also observed in the type-strain *S. wolfei* ssp. *wolfei* Göttingen DSM 2245B (27). The large number of homologous β-oxidizing genes may afford *S.* BUT1 flexibility to metabolize multiple fatty acid substrates, as its genomic population was detected in heavy ^13^C DNA during SIP with both butyrate (C_4_) and oleate (C_18_) (12). In contrast, *S. wolfei* ssp. *wolfei* is only known to β-oxidize fatty acids with four to eight carbons (28). The different homologous β-oxidizing genes may also perform identical reactions but have different kinetics and/or affinities, which could allow *S.* BUT1 to adapt to varying substrate concentrations. This is supported by the detection of the *S.* BUT1 population in ^13^C DNA in the pulse-fed digester in the previous study by Ziels *et al.* (12), but not in the continuous-fed digester in that study. Fluctuating environments are thought to lead to robustness towards gene loss within metabolic networks through an increase in multifunctional enzymes (29). Thus, the presence of various homologous genes for β-oxidation in *S.* BUT1 could have been selected for by the fluctuating environmental conditions imposed from pulse-feeding the anaerobic digester. It could also be possible that the *S.* BUT1 population was enriched in ^13^C from labelled oleate due to cross-feeding of shorter-chain intermediates during β-oxidation of the C_18_ LCFA, as other syntrophic bacteria were enriched to a high degree during growth on labelled oleate (12). Yet, the enrichment of *S.* BUT1 on ^13^C-butyrate, along with the presence of the complete butyrate β-oxidation pathway, strongly suggests that it is at least capable of β-oxidizing shorter chain fatty acids (e.g. C_4_) produced in anaerobic environments.

*Syntrophomonas* BUT1 lacks genes for aerobic or anaerobic respiration, which is similar to genomes of *S. wolfei* and *Syntrophus aciditrophicus* that are capable of syntrophic butyrate degradation (27, 30). Electrons derived from butyrate oxidation (reduced ETF from butyryl-CoA oxidation and NADH from 3-hydroxybutyryl-CoA oxidation) must be disposed through reduction of CO_2_ to formate and H^+^ to H_2_ via formate dehydrogenases and hydrogenases respectively (31–34). In the *S.* BUT1 genome, we identified genes encoding for butyryl-CoA dehydrogenase, EtfAB, and two EtfAB:quinone oxidoreductases (Supplemental File 2), indicating that this organism may transfer electrons from butyryl-CoA oxidation into membrane electron carriers using ETF. The *S.* BUT1 genome contains five gene clusters encoding for formate dehydrogenases, and four gene clusters encoding for hydrogenases (Supplemental File 2). These included a membrane-bound cytochrome b-dependent selenocysteine-containing formate dehydrogenase and [NiFe] hydrogenase that could receive butyrate-derived electrons via menaquinol (31). The quinone-binding site of the selenocysteine-containing formate dehydrogenase was on the cytoplasmic side, indicating that it likely utilizes proton motive force to drive unfavorable electron transfer to CO_2_-reducing formate generation outside of the cell. Energy investment via “reverse electron transport” is critical to drive the uphill electron transfer from butyryl-CoA/crotonyl-CoA couple to CO_2_/formate or H^+^/H_2_ couples. In contrast, the quinone binding site of the [NiFe] hydrogenase was on the periplasmic side, indicating it couples outward vectorial proton transport with H_2_ generation. Previous genomic and proteomic studies also highlight the importance of ETF-based electron transfer, membrane-bound formate dehydrogenases/hydrogenases, and reverse electron transport (6, 27, 34–37).

To complete syntrophic butyrate oxidation, NAD^+^ must also be regenerated through oxidation of NADH. However, NADH oxidation coupled with CO_2_/H^+^-reducing formate/H_2_ generation is thermodynamically unfavorable. To address this obstacle, anaerobic organisms are known to utilize electron bifurcation (or confurcation), which involves the coupling of endergonic and exergonic redox reactions to circumvent energetic barriers (38). For instance, *Thermotoga ^maritima^* utilizes a trimeric hydrogenase to couple the endergonic production of H_2_ from NADH with the exergonic production of H_2_ from reduced ferredoxin (39). Two trimeric formate dehydrogenase-and two trimeric [FeFe]hydrogenase-encoding gene clusters in *S.* BUT1 appear linked to NADH, as they all contained a NADH:acceptor oxidoreductase subunit (Supplemental File 2). Yet, if the trimeric hydrogenases and formate dehydrogenases in *S.* BUT1 produce H_2_/formate via electron bifurcation with NADH and ferredoxin, it remains unknown how *S.* BUT1 would regenerate reduced ferredoxin, as the known butyrate β-oxidation pathway does not generate reduced ferredoxin (31). Moreover, the *S.* BUT1 genome does not encode for a Rnf complex that would be necessary to generate reduced ferredoxin from NADH. Recently, the Fix (homologous to ETF) system was shown to perform electron-bifurcation to oxidize NADH coupled to the reduction of ferredoxin and ubiquinone during N_2_ fixation by *Azotobacter vinelandii* (40). The *S.* BUT1 genome encoded for a Fix-related ETF-dehydrogenase, *fixC*, as well as its associated ferredoxin, *fixX* (Supplemental File 2). A Fix system has also been detected in *S. wolfei*, and was postulated to serve as a means of generating reduced ferredoxin for H_2_ or formate production via the bifurcation mechanism (31). Yet, reduced ferredoxin production with the Fix system would be energetically costly, especially with regards to the low energy yields during syntrophic butyrate oxidation (41). Another mechanism was proposed for generating reduced ferredoxin in Rnf-lacking syntrophs that involves a heterodisulfide reductase complex (HdrABC) and ion-translocating flavin oxidoreductase genes (Flx or Flox) (42). The *flxABCD*-*hdrABC* gene cluster was shown to be widespread among anaerobic bacteria, and the protein cluster (FlxABCD-HdrABC) is proposed to function similar to the HdrABC-MvhADG cluster involved in flavin based electron bifurcation in hydrogenotrophic methanogenic archaea that couples the exergonic reduction of CoM-S-S-CoB heterodisulfide with the endergonic reduction of ferredoxin with H_2_ (43). A full *flxABCD*-*hdrABC* gene cluster was detected in the genome of *S.* BUT1 (Supplemental File 2). During the syntrophic growth of *S.* BUT1 on butyrate, the FlxABCD-HdrABC protein cluster could oxidize NADH with reduction of ferredoxin along with the reduction of a high-redox-potential disulfide acceptor (43). In *Desulfovibrio vulgaris*, it has been proposed that the DsrC serves as the high-redox thiol–disulfide electron carrier that is reduced by the FlxABCD-HdrABC complex during growth (44). The DsrC protein was also detected in the syntrophic benzoate-degrading *Syntrophorhabdus aromaticivorans* strain UI along with a *flxABCD*-*hdrABC* gene cluster (42), suggesting that the reduction of a thiol–disulfide electron carrier may be a conserved mechanism for generating reduced ferredoxin in syntrophic bacteria. Yet, the *S.* BUT1 genome does not encode for a DsrC protein, and thus an alternative and unknown thiol–disulfide electron carrier would be needed. Another possibility is that the trimeric hydrogenase can drive NADH-dependent H_2_ generation as shown in *S. wolfei* Goettingen (41). Nonetheless, this genomic analysis demonstrates that *S.* BUT1 has the potential capacity to overcome energetic barriers during syntrophic butyrate β-oxidation, and contains multiple possible mechanisms for H_2_ and formate production.

In addition to interspecies electron transfer via molecular hydrogen and formate, a potential mechanism has been proposed for direct interspecies electron transfer (DIET) in which electrons are shared via electrically-conductive nanowires (45). DIET activity has been suggested in enrichment communities degrading propionate and butyrate, in which *Syntrophomonas* was detected (46, 47). However, DIET has not been demonstrated with pure cultures of *Syntrophomonas* to date. The direct transfer of electrons is thought to depend on electrically conductive type IV pili and external polyheme cytochromes (48, 49). The *S.* BUT1 genome encodes for a type IV pilin assembly protein, *PilC*, but no genes were found that encoded for the structural protein *PilA* that is associated with DIET (49). Moreover, the type IV pilin genes identified in the *S.* BUT1 genome were of the type *Flp* (fimbrial low-molecular protein weight), which are smaller than the *Pil* type pilin utilized for DIET in *Geobacter* (50, 51). A multiheme c-type cytochrome was detected in the *S.* BUT1 genome that had 59% amino acid identity (89% coverage) with the multiheme c-type cytochrome, OmcS, from *G. sulfurreducens* that has been implicated in DIET (49) (Supplemental File 2). However, that gene also had higher homology (69% identity, 94% coverage) with the cytochrome C nitrite reductase from *S. wolfei* (accession no. WP_081424886). Therefore, the roles of DIET in the metabolism of *S.* BUT1 remain unclear, but warrant further attention via expression-based profiling.

In addition to potential genetic mechanisms for energy conservation during syntrophic growth, *S.* BUT1 also encoded for a capsule biosynthesis protein (*CapA*), which appears to be specific to syntrophic growth (52). The function of *CapA* in syntrophic growth is unclear, but may be related to the production of exopolymeric substances that facilitate interaction with methanogenic partners (52). The *S.* BUT1 genome also encoded for the *FtsW* gene that is related to shape determination, and is also a postulated biomarker of a syntrophic lifestyle (52). Based on the presence of these ‘syntrophic biomarkers’ along with genes for β-oxidization and H_2_/formate production, the genomic repertoire of *S.* BUT1 aligns with that of a syntrophic butyrate degrader.

The genome of *S.* BUT1 was compared with published genomes of the *Syntrophomonas* genus (*S*. *wolfei* subsp. wolfei, *S. wolfei* subsp. methylbutyratica, and *S*. *zehnderi*) to investigate whether metabolic genes for beta-oxidation and energy conservation were conserved (Supplemental File S4). A cutoff of 42% amino acid similarity and 80% sequence overlap was employed based on the lowest first quartile amino acid similarity we observed for top blast hits (minimum of 20% amino acid similarity and 80% overlap) of *S.* BUT1 genes to each aforementioned *Syntrophomonas* genome (42.0%, 43.5%, and 43.5%, respectively). Based on these similarity thresholds, only 34% (1050 out of 3066) of protein-coding genes in the *S.* BUT1 genome have closely related homologs present in all of the other sequenced *Syntrophomonas* genomes. Notably, 40% of the *S.* BUT1 protein-coding genes have no homologs in other *Syntrophomonas* genomes that meet the similarity criteria above. Reflecting this genomic diversity, *S.* BUT1 encodes several beta oxidation-related genes that have no homologs in the other *Syntrophomonas* genomes that meet the above criteria: one acetyl-CoA acetyltransferase, acyl-CoA dehydrogenase, acrylyl-coa reductase, and acyl-CoA thioesterase (Supplemental File S4). In addition, the *S.* BUT1 genome harbors putative isobutyryl-CoA mutase genes (SYNMBUT1_v1_1780025 – 27) highly similar to those of *Syntrophothermus lipocalidus* (65.0-83.4% amino acid similarity), suggesting that *S.* BUT1 may also be capable of syntrophic isobutyrate degradation. Hydrogenases, formate dehydrogenases, and energy conservation genes were generally conserved among *S.* BUT1 and the other *Syntrophomonas* genomes. Only the cytochrome b-dependent [NiFe] hydrogenase has no homologs in the *S. wolfei* subsp. *wolfei* genome. This implies that *S.* BUT1 may have distinct capabilities for fatty acid oxidation, but the energy conservation necessary to drive syntrophic beta oxidation may not vary between *Syntrophomonas* species.

A genomic analysis of the *Methanothrix* BUT2 genome indicated that it contained the complete pathway for methane production from acetate (Figure 4, Supplemental File 3). This observation agrees with the physiology of other *Methanothrix* species, which are known aceticlastic methanogens (53, 54). *M.* BUT2 also contained genes that likely are involved in energy conservation during aceticlastic methanogenesis. The genome of *M.* BUT2 harbored acetyl-CoA synthetase for acetate activation, bifunctional CO dehydrogenase/acetyl-CoA synthase (CODH/ACS) to oxidatively split acetyl-CoA into CO_2_ and CH_3_-H_4_MPT, tetrahydromethanopterin S-methyltransferase, and methyl-CoM reductase for methyl-CoM reduction to CH_4_ (Supplemental File 3). To couple acetyl-CoA oxidation and reductive CH_4_ generation, BUT2 must transfer electrons from Fd_red_ to CoM-SH/CoB-SH. We identified a FpoF-lacking F_420_H_2_ dehydrogenase (Fpo) complex and heterodisulfide reductase (HdrDE) that could facilitate this (Supplemental File 3) and also generate an ion motive force (55). This energy conservation system is highly similar to *M. thermophila* acetate oxidation (55). In previous studies, *Methanothrix* species have been observed to co-occur with *Syntrophomonas* in LCFA-degrading (56) and butyrate-degrading (57–59) anaerobic environments. In this study, the stable isotope-informed metagenomic analysis strongly suggests that the labelling of *M.* BUT2 DNA was due to the incorporation of ^13^C-acetate produced during the degradation of ^13^C-butyrate by *S.* BUT1.

A near-complete pathway for methane production from CO_2_ was also observed in the *M.* BUT2 genome (Supplemental File 3). The only gene lacking in the CO_2_-reducing pathway was a F_420_-dependent N_5_N_10_-methylene-tetrahydromethanopterin dehydrogenase (Mtd). While *Methanothrix* are thought to be obligate aceticlastic methanogens (53, 54), the presence and expression of the CO_2_-reducing pathway in *Methanothrix* was previously reported (60–62) and was hypothesized to be involved in methane formation via DIET. However, the mechanism through which *Methanothrix* would directly accept electrons from its syntrophic partner has not been identified (60, 61). The other known electron donors for methane production from CO_2_ are hydrogen and formate. A membrane bound hydrogenase (*mbhAB*) was observed in the *M.* BUT2 genome (Supplemental File 3). In other studies, negligible hydrogenase activity was observed with *Methanothrix* species (63). Two monomeric formate dehydrogenase enzymes (*fdhA*) were also encoded in *M.* BUT2 (Supplemental File 3). Experiments with thermophilic *M.* sp. strain CALS-1 and mesophilic *M. concilii* showed that they displayed formate dehydrogenase activity by splitting formate into hydrogen and CO_2_, however the produced CO_2_ was not used for methane generation (63, 64). Yet, the mesophilic *M. soehngenii* did not show formate dehydrogenase activity (54). Thus, the roles of the hydrogenases, formate dehydrogenases, and CO_2_-reducing pathway for methane generation in *M.* BUT2 are not clear. Transcriptomic or proteomic approaches are needed to elucidate the activity of the CO_2_-reducing methanogenesis production pathway during syntrophic growth on butyrate with *S.* BUT1.

## Conclusions

In this study, stable isotope-informed genome-resolved metagenomics was used to provide genomic insight into syntrophic metabolism during butyrate degradation in anaerobic digesters. The results obtained via genome binning and metabolic reconstruction showed that the ^13^C-enriched *Syntrophomonas* genome contained the genetic capacity to convert butyrate into precursor metabolites for methane formation—acetate, hydrogen and formate. The ^13^C-enriched *Methanothrix* genome likely consumed the acetate produced during butyrate degradation, incorporating some ^13^C into biomass. The presence of a CO_2_-reducing pathway, as well as formate dehydrogenase and hydrogenase genes, in the *Methanothrix* genome leaves open the possibility of flexible metabolism during methanogenesis. As syntrophic fatty acid degrading populations are often slow-growing and thus difficult to isolate, this study demonstrates a new approach to link ecophysiology with genomic identity in these important populations involved in anaerobic biotechnologies, as well as global carbon cycling. Advancing our understanding of *in-situ* metabolic activities within anaerobic communities is paramount, as these microbiomes contain multiple interacting functional groups that, in cooperation, enable the processing of degradable organic carbon into methane gas. Coupling SIP-informed metagenomics with other activity-based techniques, such as metabolomics, transcriptomics, and proteomics, could further illuminate the structure of anaerobic metabolic networks as well as quantify metabolite fluxes, thus enabling newly informed process models to predict rates of anaerobic carbon transformation.

## Experimental Procedures

### Batch incubations with ^13^C-labelled butyrate

Two 4 L anaerobic digesters treating dairy manure and sodium oleate were operated for over 200 d at a solids retention time of 20 d and a temperature of 35°C, as described by Ziels *et al.* (56). The two digesters were operated with different feeding frequencies of sodium oleate. One digester received sodium oleate once every 48 hrs, while the other digester was fed semi-continuously every 6 hrs (56).

On day 228 of digester operation, 10 mL samples were collected from each digester, and immediately transferred to 35 mL glass serum bottles that were pre-purged with N_2_:CO_2_ (80:20), and capped with butyl rubber septa. Duplicate microcosms were fed with a 1 M solution of either ^12^C sodium butyrate or ^13^C-labeled sodium butyrate (>98% atom purity, Cambridge Isotope Laboratories, Tewksbury, MA, USA) to reach an initial butyrate concentration of 40 mM. The ^13^C-labeled sodium butyrate was universally labeled at all 4 carbons. Triplicate blank controls were incubated in parallel to measure background methane production from the inoculum. Methane production was measured approximately every 4 hr over the 50 hr incubation time using a digital manometer (Series 490 A, Dwyer Instruments) and GC-FID (SRI 8610C), according to Ziels *et al.* (56).

### Stable isotope probing

DNA was extracted from the duplicate 10 mL microcosms after the 50-hr incubation, and was separated via density-gradient centrifugation and fractionated as previously described (12). DNA was measured in 24 density gradient fractions using QuBit (Invitrogen, MA, USA). *Syntrophomonas* 16S rRNA genes were quantified in gradient fractions as described by Ziels et al. (12), using previously developed primers and probes (65). Heavy DNA fractions with buoyant densities between 1.70-1.705 g/mL (Supplemental Figure S2) were selected for each microcosm sample and sent for metagenomic sequencing at MR DNA Laboratories (Shallowater, TX, USA), as well as for 16S rRNA gene iTag sequencing at the U.S. Department of Energy Joint Genome Institute (JGI) according to Ziels *et al.* (12). Metagenome libraries were prepared using the Nextera DNA sample preparation kit (Illumina Inc., Hayward, CA, USA) following the manufacturer’s instructions. The metagenome libraries were sequenced in 150 bp paired-end mode on a HiSeq 2500 (Illumina Inc., Hayward, California, USA). Bioinformatic analysis of the 16S rRNA iTags is described in detail in the Supporting Information.

### Genome binning, annotation, and statistical analysis

All metagenomic reads were initially trimmed and quality filtered using illumina-utils (66) (available from: https://github.com/merenlab/illumina-utils) according to the parameters of Minoche et al. (67). Metagenomic reads from all ^13^C-butyrate fed microcosms were co-assembled using MEGAHIT v1.1.1 (68). Open reading frames were called with Prodigal v.2.6.3 (69), and were taxonomically classified with GhostKOALA (70). Short reads from the ^12^C and ^13^C metagenomes were mapped onto the contigs using Bowtie2 (71) with default parameters. Additionally, metagenomic reads from the total biomass collected from each digester 2 days after the butyrate SIP experiment (i.e. time zero) (12) were mapped onto the assembled contigs to facilitate the subsequent differential coverage binning. The contigs were then binned according to the workflow of Eren *et al.* (72) using Anvi’o v.2.4.0, as described in detail in the Supplemental Information. After manual refinement of the bins using Anvi’o, we obtained a set of 160 genomic bins that were assessed for completeness and contamination with CheckM (72) (Supplemental File 1). Differential abundance of each genomic bin in the ^13^C and ^12^C butyrate metagenomes of each digester was determined using DESeq2 (14) using mapped read counts. A significant difference in abundance between ^12^C and ^13^C metagenomes was established by a *p* value less than 0.05. The average nucleotide identity (ANI) between ^13^C-enriched genomic bins and publicly-available genomes from closely-related organisms were calculated with pyANI (available from: https://github.com/widdowquinn/pyani). Open reading frames were annotated with the MicroScope platform (73), and metabolic reconstructions were performed in Pathway Tools (74). Potential type IV pilin genes were identified with the PilFind program (50).

### Data Availability

We have made publicly available the following: raw sequence reads and metagenome assemblies for the butyrate DNA-SIP metagenomes under NCBI Sequence Read Archive under BioProject PRJNA524401; genomic FASTA files for each ^13^C-enriched genomic bin (https://doi.org/10.6084/m9.figshare.7761776); and the annotation data for the two ^13^C enriched MAGs (https://doi.org/10.6084/m9.figshare.7761710). The time-zero raw metagenomic reads from the study by Ziels et al. (12) that were used for differential coverage binning, are available via the U.S. Joint Genome Institute Genome Portal (https://genome.jgi.doe.gov/portal/) under Project IDs: 1105507, 1105497. 16S rRNA gene amplicon sequences are available via the U.S. Joint Genome Institute Genome Portal under Project Number: 1105527, Sample IDs: 112232-112239.

## Acknowledgments

This work was funded by US EPA STAR [grant RD835567] and ERC [grant 323009]. The work conducted by the U.S. Department of Energy Joint Genome Institute, a DOE Office of Science User Facility, is supported by the Office of Science of the U.S. Department of Energy under Contract No. DE-AC02-05CH11231.We acknowledge H. David Stensel and Alfons Stams for their helpful input.

