## Supplemental Information for "Elucidating syntrophic butyrate-degrading populations in anaerobic digesters using stable isotope-informed genome-resolved metagenomics"

**Supplemental Methods:**

*16S rRNA iTag Sequence Analysis*

Raw reads were filtered by trimming the first 10 bp, truncating forward reads at 265 bp, truncating reverse reads at 180 bp, and filtering all reads based on a maximum expected error of 2 using DADA2 (Callahan *et al.*, 2016). The filtered and trimmed reads were then dereplicated and denoised into exact sequences using estimated error parameters with DADA2. Forward and reverse sequences were then merged with DADA2 using a minimum overlap of 20 bp and zero allowed mismatches. Merged and denoised sequences were then truncated to 390 bp and clustered into OTUs with a 99.5% similarity cutoff following chimera removal with UPARSE v.8.1 (Edgar, 2013). Representative sequences of the 99.5% OTUs were classified against the SILVA SSU Ref NR dataset v.123 using the RDP classifier (Wang *et al.*, 2007).

*Binning metagenome-assembled genomes (MAGs) with Anvi’o*

After co-assembly, read mapping, and open reading frame prediction as described in the manuscript, metagenomic contigs were binned using Anvi’o v.2.4.0. Single-copy genes were searched using the “anvi-run-hmms” command. The “anvi-profile” command was used to parse contig coverage across all samples from the BAM files with samtools (Li *et al.*, 2009). The “anvi-merge” command was used to compile the coverage information for contigs across all samples into single anvi’o profile. The initial binning was conducted with the “anvi-cluster-with-concoct”, which uses CONCOCT (Alneberg *et al.*, 2014), by constraining the number of bins to 40 (“--num-clusters 40”) to minimize fragmentation error (i.e. splitting up a single bin into multiple smaller bins) (Lee *et al.*, 2017). Bins that displayed “conflation error” (i.e. a bin has multiple populations and/or contamination) (Lee *et al.*, 2017) were interactively refined the binned contigs using the “anvi-refine” command based on completion and redundancy estimates from the presence of bacterial and archaeal single-copy genes, taxonomy of ORFs from BlastKOALA, tetra-nucleotide frequency, and coverage patterns across multiple samples.

**Supplemental Figures**

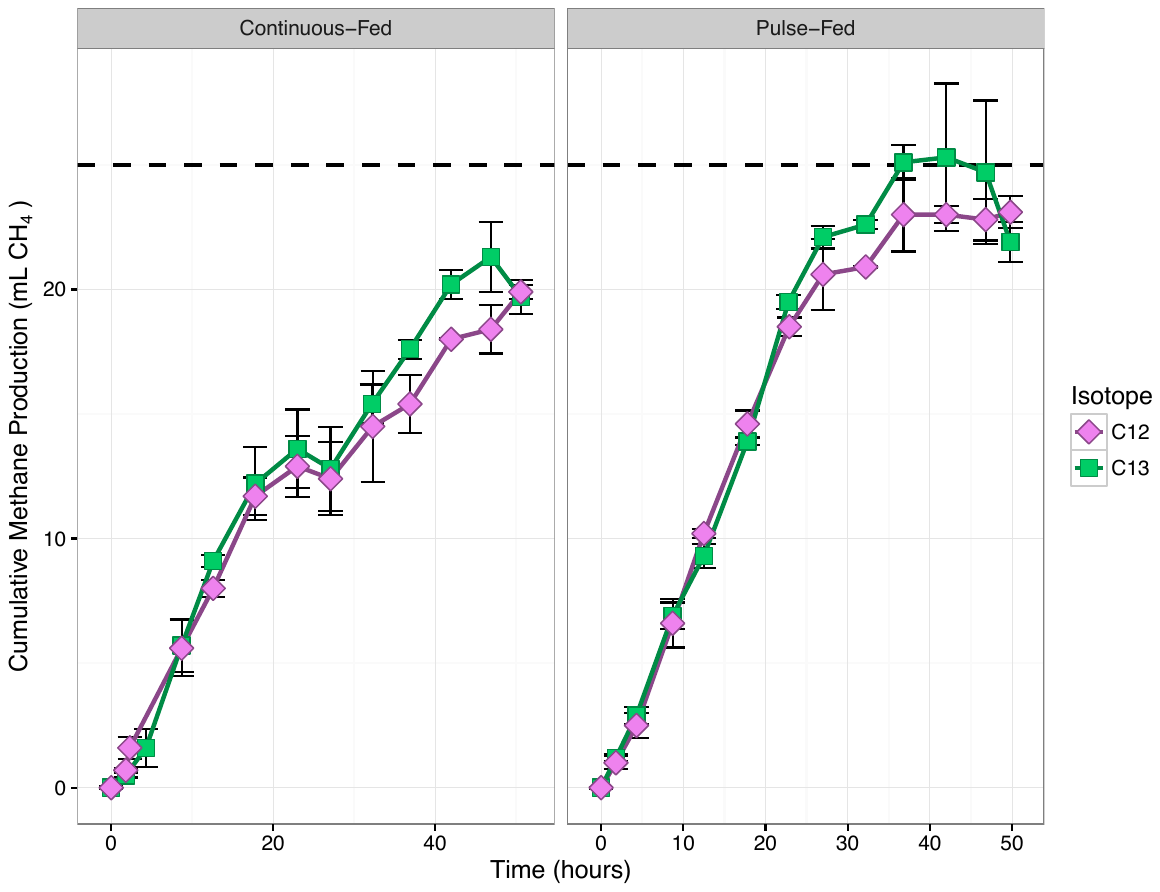

**Supplemental Figure S1:** Cumulative methane production (minus blank controls) for the microcosms fed with ^12^C- and ^13^C-labeled butyrate over approximately 50 hours. The black dashed line shows the theoretical methane potential of the added oleate (25.3 ml CH_4_; based on 1.82 g COD/g butyrate, 40 mM concentration, 10 ml sample, and 35 °C temperature). Error bars represent the standard deviation of the biological replicates.

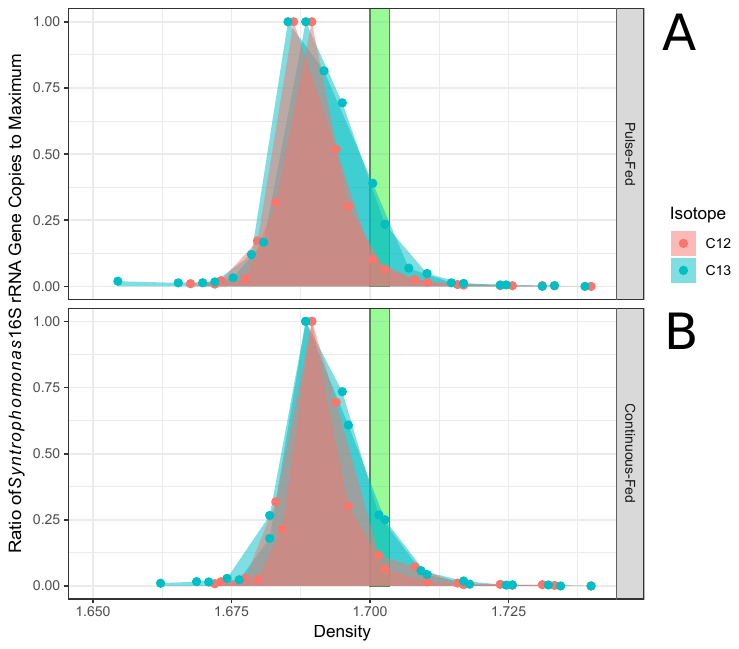

**Supplemental Figure S2:** Ratios of *Syntrophomonas* 16S rRNA genes measured by qPCR in each density-gradient fraction to the maximum observed across all density fractions. Fractions were recovered from isopycnic separation of DNA from ^13^C-incubated microcosms and ^12^C-controls for both anaerobic digesters. The green rectangles indicate density gradient fractions that were pooled for subsequent 16S rRNA gene amplicon sequencing and metagenomic sequencing.

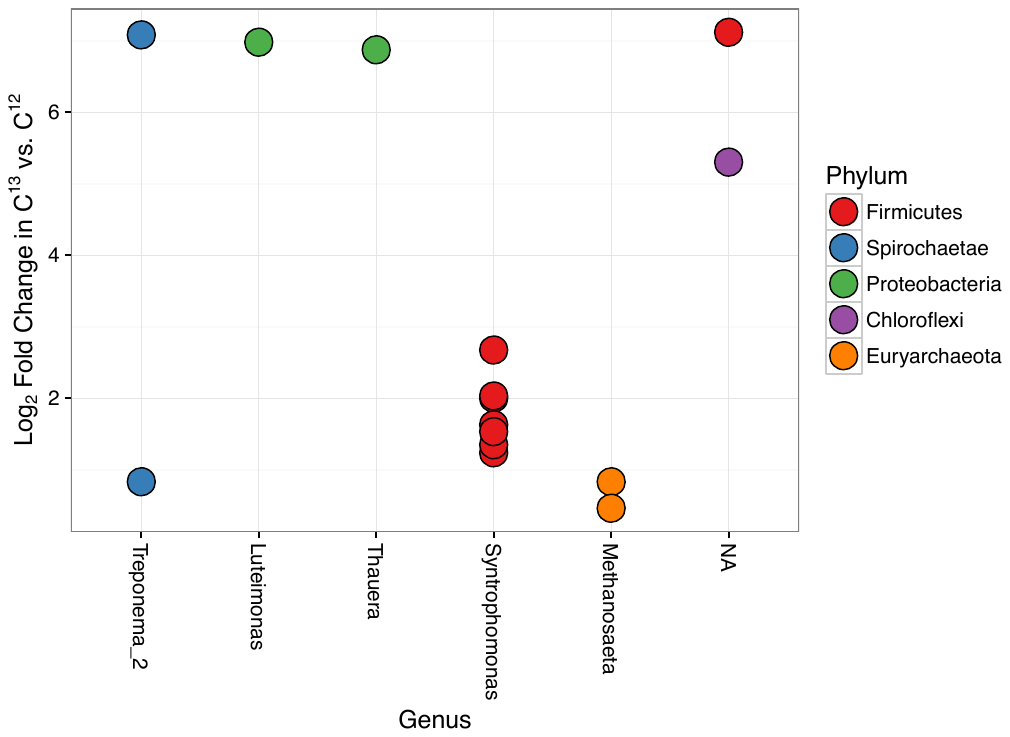

**Supplemental Figure S3:** Genus and phylum level taxonomic assignments of 16S rRNA amplicon OTUs in the pulse-fed codigester that were identified as significantly enriched in ^13^C samples versus ^12^C samples using DESeq2 (Love et al., 2014), along with their log_2_ fold change.

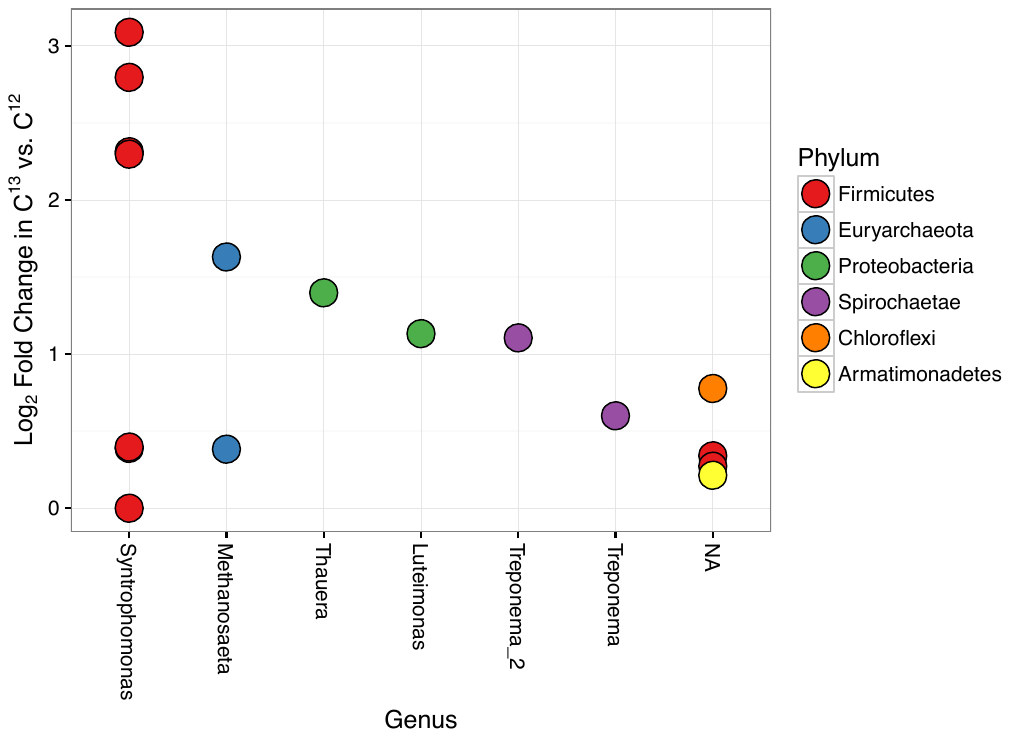

**Supplemental Figure S4:** Genus and phylum level taxonomic assignments of 16S rRNA amplicon OTUs in the continuous-fed codigester that were identified as significantly enriched in ^13^C samples versus ^12^C samples using DESeq2 (Love et al., 2014), along with their log_2_ fold change.

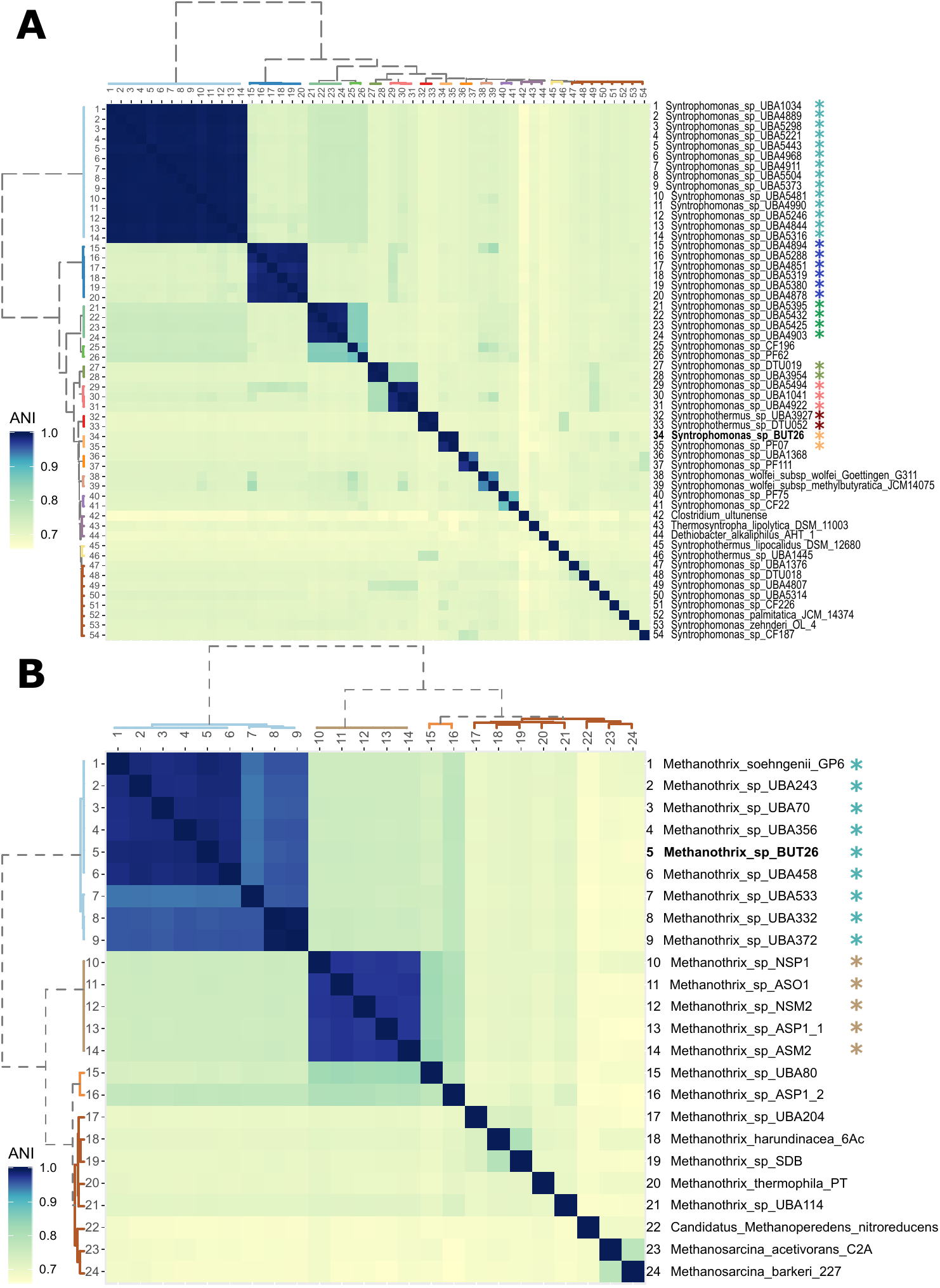

**Supplemental Figure S5:** Heatmaps of average nucleotide identity (ANI) between genomes from (A) *Syntrophomonadaceae* and (B) *Methanosarcinales*. Genomes were clustered based on the ANI values using Ward’s minimum variance method. The genome names shown in bold were identified in this study. Other genomes were obtained via the NCBI nr database (downloaded April, 2018)

**Supplemental Tables**

**Supplemental Table S1:** Summary of SIP metagenomes utilized for the co-assembly, including number of raw and filtered reads, and the fraction of reads that mapped to co-assembly.

| **Digester** | **Replicate** | **Isotope** | **Raw Reads** | **Filtered Reads** | **Reads Mapped to Co-assembly (%)** |
| --- | --- | --- | --- | --- | --- |
| Pulse Fed | A | ^12^C | 31,425,432 | 28,097,374 | 63.98 |
| Pulse Fed | B | ^12^C | 31,646,858 | 28,395,122 | 64.14 |
| Pulse Fed | A | ^13^C | 40,852,542 | 36,232,668 | 68.16 |
| Pulse Fed | B | ^13^C | 25,672,716 | 23,368,004 | 69.12 |
| Continuous Fed | A | ^12^C | 36,841,972 | 32,975,712 | 69.97 |
| Continuous Fed | B | ^12^C | 35,962,594 | 32,641,070 | 66.65 |
| Continuous Fed | A | ^13^C | 29,134,756 | 26,823,396 | 69.34 |
| Continuous Fed | B | ^13^C | 34,718,070 | 31,132,802 | 70.26 |
